# Sustained Ca^2+^ mobilizations: a quantitative approach to predict their importance in cell-cell communication and wound healing

**DOI:** 10.1101/558320

**Authors:** Yoonjoo Lee, Min Tae Kim, Garrett Rhodes, Kelsey Sack, Sung Jun Son, Celeste B. Rich, Vijaya B. Kolachalama, Christopher V Gabel, Vickery Trinkaus-Randall

## Abstract

Epithelial wound healing requires the coordination of cells to migrate as a unit over the basement membrane after injury. To understand the process of this coordinated movement, it is critical to study the dynamics of cell-cell communication. We developed a method to characterize the injury-induced sustained Ca^2+^ mobilizations that travel between cells for periods of time up to several hours. These events of communication are concentrated along the wound edge and are reduced in cells further away from the wound. Our goal was to delineate the role and contribution of these sustained mobilizations and using MATLAB analyses, we determined the probability of cell-cell communication events in in vitro models and ex vivo organ culture models. We demonstrated that the injury response was complex and represented the activation of a number of receptors. In addition, we found that pannexin channels mediated the cell-cell communication and motility. Furthermore, the sustained Ca^2+^ mobilizations are associated with changes in cell morphology and motility during wound healing. The results demonstrate that both purinoreceptors and pannexins regulate the sustained Ca^2+^ mobilization necessary for cell-cell communication in wound healing.

## Introduction

The epithelium serves as a barrier to external disruptions such as injury or environmental factors and repair requires coordination between cells to migrate over the basement membrane and close the wound. To understand how epithelial cells move as a unit after injury, the dynamics of cell-cell communication and coordination of the process need to be studied. An excellent model tissue is the corneal epithelium, which is an avascular stratified squamous tissue that responds to growth factors and nucleotides when the epithelial barrier is damaged. One signal that has a ubiquitous response in epithelial wound healing is the release of the nucleotide, ATP, which may occur because of a change in cell shape as in bronchial epithelia and corneal epithelial injury, or an alteration in force such as in glaucoma [1-3]. Within milliseconds to seconds after injury, extracellular ATP binds to purinoreceptors and triggers a transient Ca^2+^ wave, which is used by cells to transduce mechanical signals into chemical signals and alter signaling pathways [4-8]. This response is mimicked in unwounded cultures that are exposed to medium collected from injured cells; however the response is absent when the wound medium is pretreated with apyrase, an ectonucleotidase [5].

Nucleotides are ligands for a number of purinergic receptors, such as P2Y [G protein-coupled receptors (GPCR)] and P2X (ligand gated ion channels), which are known to mediate cell migration, proliferation, and inflammation [9]. Nucleotides affect the phosphorylation of a number of proteins, including epidermal growth factor receptor (EGFR), Src, extracellular-signal-regulated protein kinase (ERK), β4 integrin, and paxillin [5,7,10]. Furthermore knocking down the G-protein purinoreceptor, P2Y2, results in a decrease in Ca^2+^ mobilizations, wound healing and phosphorylation of paxillin (Y118) and EGFR (Y1068), but not EGFR (Y1173) [7,11]. In contrast knocking down the P2Y4 receptor did not significantly reduce the injury-induced response [11]. In addition, only the P2Y2 receptor increased in expression after injury [7]. Another purinoreceptor where significant changes were detected was P2X7, which exhibited a planar polarity after injury, was prominent at the leading edge and where the polarity was abrogated in the pre-diabetic model [10]. Furthermore, P2X7 expression was significantly elevated in diabetic corneas, and the corneal epithelium of a murine pre-diabetic model displayed a similar elevated mRNA [12-13]. These data led us to hypothesize that the purinoreceptors act as sensors. Additional evidence from other cell systems suggests the presence of a feed-forward system where ATP moves through pannexin channels and activates P2X7 receptors [14]. This type of system would suggest a continuous release of ATP along the wound margin. For example, ATP is released by neutrophils and appears to act for chemotaxis during inflammation while in another cell system, the channel protein, pannexin1, plays a role in cell migration during injury in dendritic cells [15-18].

In this study we developed a novel method to identify and characterize the degree of cell-cell communication that occurs through sustained Ca^2+^ mobilizations after injury, which are concentrated along the epithelial wound edge and reduced in cells distal to the injury. Using MATLAB analyses, we generated profiles of the sustained Ca^2+^ mobilizations, and demonstrated that the Ca^2+^ response was replicated in ex vivo organ culture models. The sustained Ca^2+^ mobilizations were present also after stimulation with either BzATP or UTP, and the probability that cells would communicate was greater in response to BzATP. The specificity was demonstrated using competitive inhibitors of P2Y2 and P2X7, AR-C 118925XX and A438079 respectively. Likewise, an inhibitor of pannexin1 attenuated both the wound and BzATP agonist initiated response. These sustained mobilizations are correlated with changes in cellular morphology and motility, which were prominent in cells at the leading edge during cell migration after wounding. Together, our results demonstrate that the sustained Ca^2+^ mobilizations mediated by purinoreceptors and pannexins are a vital component in regulating the long-term response to injury.

## Materials and Methods

### Reagents

ATP, 2,3-O-(4-benzoylbenzoyl)-ATP (BzATP), and UTP were all purchased from Sigma-Aldrich (St. Louis, MO). A438079 hydrochloride, AR-C 118925XX, 10Panx Inhibitory peptide, and scramble control peptide were purchased from Tocris Biosciences (Minneapolis, MN). Pannexin-1 polyclonal rabbit antibodies were purchased from Alomone (Jerusalem, Israel), and Connexin-43 (Cx43) polyclonal rabbit antibodies were purchased from Santa Cruz Biotechnology (Santa Cruz, CA).

### Cell culture

Human corneal limbal epithelial (HCLE) cells, a gift from Dr. Gipson (Schepens Eye Research Institute/Mass. Eye and Ear, Boston, MA) were evaluated for mycoplasm [19]. The HCLE cell line was verified at Johns Hopkins DNA Services (Baltimore, MD). Cells were maintained in Keratinocyte Serum-Free Media (KSFM) with growth supplements (25-μg/mL bovine pituitary extract, 0.02 nM EGF, and 0.3 mM CaCl_2_). Cells were passaged when 70-80% confluent and plated on either glass bottom dishes (MatTek Corporation, Ashland, MA) for live cell imaging, scratch wound assays, and immunofluorescence, or on cell culture-treated plastic petri dishes for Western blot analysis for approximately 72 hours prior to experimentation at a density of 150 cells/mm^2^. Approximately 16-24 hours before experimentation, the media was changed to unsupplemented KSFM, as previously described [10].

### Organ culture and tissue preparation

The research protocol conformed to the standards of the Association for Research in Vision and Ophthalmology for the Use of Animals in Ophthalmic Care and Vision Research and the Boston University IACUC. C57BL/6J mice were obtained from Jackson Laboratory (The Jackson Laboratory; Bar Harbor, ME). For organ culture live imaging, the corneas were enucleated and incubated in KSFM at 37 °C and 5% CO_2_. To prepare tissues for immunohistochemistry, a 1.5mm-diameter trephine was used to delineate the region in the central cornea that would be wounded by removing or abrading the epithelium. After wounding, the corneas were dissected, leaving an intact scleral rim, and incubated in Dulbecco’s modified Eagle’s medium (DMEM) at 37 °C and 5% CO_2_, as described [10, 20].

### Live cell confocal imaging

#### Ca^2+^ mobilization studies

All image studies were imaged on the Zeiss Axiovert LSM 880 confocal microscope, with the ex vivo live-imaging utilizing the FAST module and AIRYScan (Zeiss, Thornwood, NY). Ca^2+^ mobilization was performed on HCLE cells as previously described and on ex vivo mouse corneas [10, 21]. For in vitro imaging, HCLE cells were cultured to confluence on glass bottom dishes and pre-loaded with 5 μM Fluo-3AM fluorescent dye (Invitrogen, Carlsbad, CA) to allow for Ca^2+^ visualization, at a final concentration of 1% (v/v) DMSO and 2% (w/v) pluronic acid at 37°C and 5% CO_2_ [10]. Images were collected after any of the following experiments: agonist stimulation by addition of either BzATP or UTP (final concentration of 25 μM), or scratch-wound injury and taken every 3 seconds for up to 2 hours in length. For ex vivo imaging of mouse corneas, the corneas were mounted on glass bottom dishes and preincubated with 50 μM Fluo-3AM fluorescent dye for one hour and CellMask™ Deep Red Plasma membrane stain, which was used at 1:10000 (CellMask™:media) (Thermo Fisher, Waltham, MA), at a final concentration of 1% (v/v) DMSO and 20% (w/v) pluronic acid for 30 minutes at 37°C and 5% CO_2_.

#### Cell shape changes and migration

To examine cell migration and alterations in cell shape, HCLE cells were pre-loaded with either CellMask™ Deep Red Plasma membrane stain as described above or 1 μM of SiR-Actin Spirochrome probe (Cytoskeleton Inc., Denver, CO) for 10 minutes at 37°C and 5% CO_2_, for imaging of F-actin. Both long- and short-term studies were performed. For long-term studies, images were collected immediately after injury and every 5 minutes for 6 hours on a Zeiss Axiovert LSM 880 confocal microscope. For short-term studies, images were taken immediately after injury and every 5 seconds for up to 2 hours on a Zeiss Axiovert LSM 880 confocal microscope. Analyses for all the described image studies were performed using FIJI/ImageJ (NIH, Bethesda, MD; http://imagej.nih.gov/ij/) along with MATLAB programs (MATLAB, MathWorks, Inc.) written for the analysis described below.

### Modeling of Ca^2+^ waves

To analyze spatiotemporal communication between individual cells or groups of cells, videos were collected from each experiment and exported in TIF or AVI format. Two different custom MATLAB scripts were employed to analyze Ca^2+^ responses based on cell population (individual cells or a population of cells). The individual cell analysis technique was previously described [22]. To examine cell-cell communication we developed a script to 1) Identify Ca^2+^ events and generate an event kymograph; and 2) Calculate the probability of neighboring cells having a Ca^2+^ event that was induced within 10 frames after an established Ca^2+^ event had occurred in a given cell. Cell positions were marked by either an automated computer program or manual detection. Signaling events within each trace were identified as being greater than a threshold of 50% of the maximum normalized fluorescent signal. A cluster was defined as a group of 2-3 adjacent cells where Ca^2+^ mobilizations occurred, and the number of clusters was measured over time. Neighboring cells were identified as all cells within <35 μm of one another. The neighboring cells displaying events within 10 frames (30 seconds) of each other were scored as “correlated” events. The probability that an event in any particular cell triggered a correlated event in any of its neighbors was calculated and defined as the “event probability.”

### ATP release assay

To determine concentration of ATP released after injury, HCLE cells were plated on culture-treated plastic and grown to confluence in KSFM contianing growth supplements. The growth supplements were removed from the media 24 hours before wounding. To wound the cells, a comb made from plastic gel-loading tips was used to make a scratch wound, and the media was collected every 20 minutes and clarified by centrifugation at 663 x g. The clarified media was collected and stored on ice until ready for analysis with a luciferase-based ATP Determination Kit (Invitrogen, Carlsbad, CA). Samples were vortexed and 5 μL aliquots were plated on a white-bottomed 96-well plate (Corning). A reaction buffer (0.5 mM D-luciferin, 1.25 μg/mL firefly luciferase, 25 mM Trycine buffer, pH 7.8, 5 mM MgSO4, 100 μM EDTA, and 1 mM DTT) was prepared immediately before analysis and protected from light. To determine ATP levels, luciferase-generated luminescence was detected using a BioTek Synergy HT plate reader with injector (BioTek, Winooski, VT). A standard curve of ATP was made in KSFM only. To ensure equal time for each reaction, 95 μL of reaction buffer was injected into a well and allowed to incubate for four seconds before luminescence was read. ATP levels were calculated from raw luminescence values using the standard curve.

### Immunofluorescence and confocal microscopy

HCLE cells and mouse corneas were fixed in freshly prepared 4% paraformaldehyde in PBS for 20 minutes at room temperature (cells) or overnight at 4°C (corneas). Immunofluorescent staining was performed [10]. Briefly, cells and corneas were permeabilized with 0.1% (v/v) Triton X-100 in PBS for 2-5 minutes and blocked with 4% BSA in PBS (blocking solution) for 1 hour. Cells and corneas were incubated in primary antibody solutions overnight at 4°C, and the following day they were incubated with the corresponding Alexa Fluor-conjugated secondary antibody (Invitrogen, Carlsbad, CA) at a dilution of 1:100 in blocking solution for 1 hour at room temperature. Rhodamine-conjugated phalloidin (Invitrogen, 1:50) was used to visualize F-actin. Cells and corneas were mounted using VectaSHIELD with DAPI (Vector Labs, Burlingame, CA). Images were obtained on a Zeiss LSM 700 (Zeiss, Thornwood, NY) confocal microscope with indicated objectives and settings, and analyzed using ZEN (Zeiss, Thornwood, NY) or FIJI/ImageJ (NIH, Bethesda, MD; http://imagej.nih.gov/ij/).

### Statistical Analysis

At least three independent experiments were run for each set of samples, and the mean ± standard error of the mean (SEM) was determined. Statistical significance was determined by unpaired, one-tailed Student’s t-test or two-way ANOVA with appropriate post hoc tests using GraphPad Prism 5 (GraphPad Software, San Diego, CA) and R studio (RStudio, Inc., Boston, MA).

## Results

### Sustained Ca^2+^ mobilizations after injury recruit cells along the wound margin

In this study, we investigated the hypothesis that sustained Ca^2+^ mobilizations are responsible for cell-cell communication, which underlies the collective cell migration of corneal epithelial cells after agonist stimulation or injury. First, we examined the Ca^2+^ mobilization within HCLE cells by live-cell imaging before and after wounding. Single-frame images of the Ca^2+^ mobilization before and after (0, 5, and 120 mins) a scratch wound are shown (Fig 1A, first panel). Immediately after wounding, there is a large mobilization of Ca^2+^ that is transient and has been described [2] (Fig 1A, first panel 0 min). To examine the response we outlined the cells along the leading edge of the wound, thereby depicting the regions of interest (ROI: Fig 1A middle panel, outlined in white). From this data, we generated a kymograph (Fig 1A, third panel), representing each of the individual leading edge cells (ticks along the y-axis) and the changes in fluorescence intensity over time (Fig 1A, third panel). At t=0 (wounding), the initial Ca^2+^ wave was observed, as indicated by the high intensity of Ca^2+^ (intensity scale) and this is followed by Ca^2+^ mobilizations that are sustained for up to two hours (Fig 1A kymograph, S1 Movie). We speculated that regions of neighboring cells at the leading edge displayed a synchronicity, indicating that the transfer of information between cells may be involved in wound healing (Fig 1A kymograph). The cells back from the leading edge, denoted by being at least two cells distal from the wound, were less active compared to the cells at the leading edge of the wound (S1 Fig).

**Fig 1.**
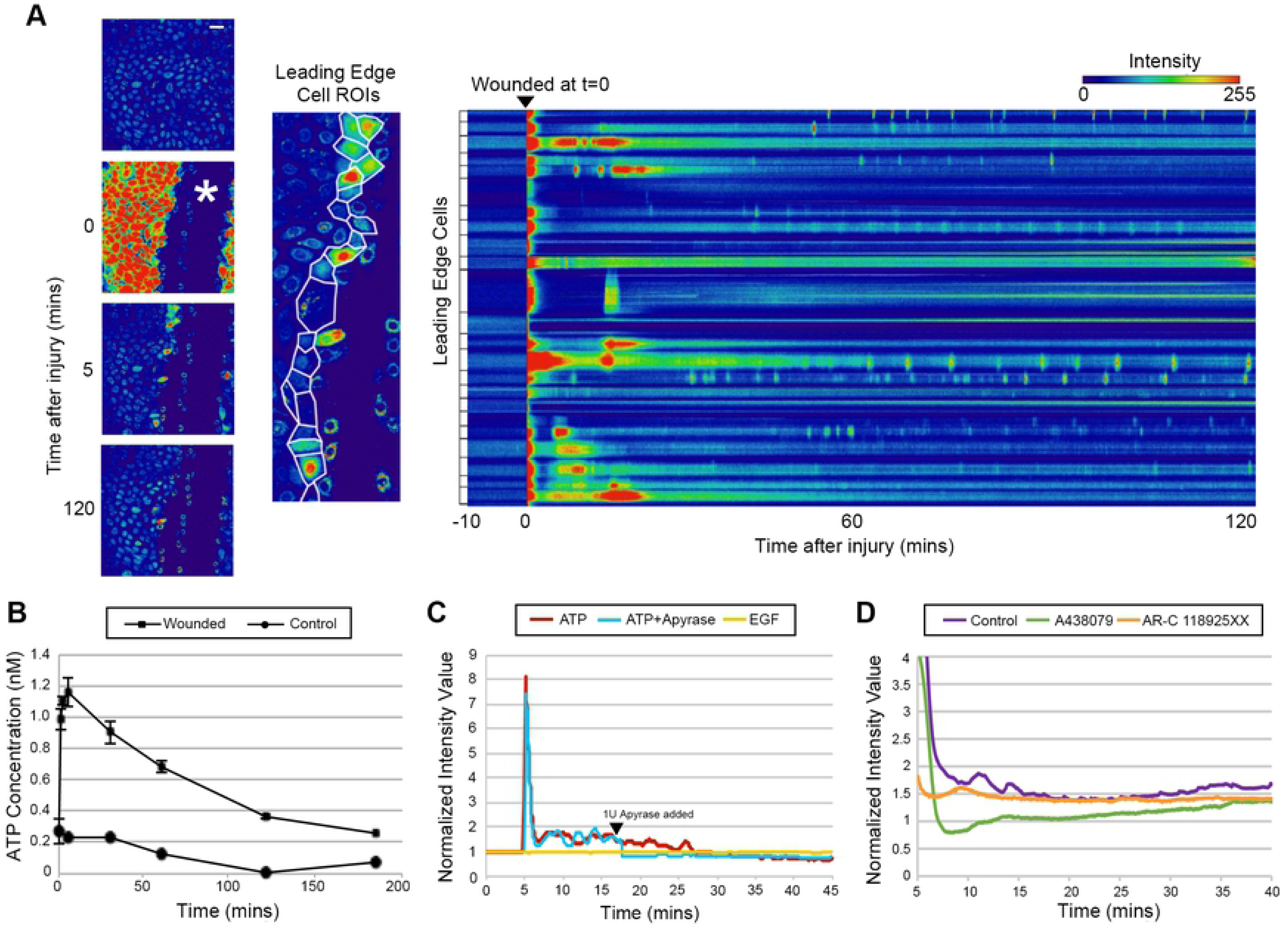
Communication between cells after injury is inhibited by purinergic inhibitors. (A) Representative images of the initial Ca^2+^ wave and the sustained Ca^2+^ mobilizations upon scratch wounding (wound marked with asterisks). Based on the outlined cells closest to the wound edge (in white), kymographs of the cells closest to the wound edge were generated to observe fluorescent intensity changes over time. The brackets on the left of each horizontal row represents the activity of a single cell. Cells were preincubated in 5 μM Fluo3-AM for 30 minutes and imaged on the Zeiss Axiovert LSM 880 confocal laser scanning microscope. Scale bar = 60 μm. (n=7). (B) Graph of the mean extracellular ATP concentration over time after injury compared to unwounded control. Error bars represent SEM (n=3). (C) Comparison of representative fluorescence over time after treatment with ATP, ATP and apyrase, and EGF. Black arrowhead marks the time apyrase was added. Apyrase abrogated the Ca^2+^ response. EGF induced negligible Ca^2+^ mobilizations compared to that of ATP. (n=4). (D) Injury induced response (Normalized intensity value of fluorescence) in presence or absence of inhibitors: A438079 (P2X7 competitive inhibitor) or AR-C 118925XX (P2Y2 competitive inhibitor) (n=4).

In addition to live-cell imaging of the Ca^2+^ mobilization, ATP in the media from HCLE cells (control and wounded) was examined by luciferase assay. As seen in Figure 1B, the concentration of ATP in wounded media was significantly greater than control for at least three hours. Moreover, it was six-fold higher in the wounded sample than the unwounded control after 60 minutes (Fig 1B). This continuous presence of extracellular ATP after injury could be responsible for maintaining the sustained Ca^2+^ mobilizations (Fig 1A).

To assess the role of ATP, we compared the response of unwounded cells to ATP, ATP + apyrase, or EGF and measured the normalized intensity value of fluorescence. Previously we had demonstrated the role of ATP and the ectonucleotidase, apyrase, on the initial mobilization [2, 5]. In the current experiments we demonstrated the role of apyrase on sustained mobilizations (Fig 1C, red and blue lines) by adding apyrase after the initial Ca^2+^ wave (arrowhead). The addition quenched the subsequent Ca^2+^ mobilizations (Fig 1C, blue line), indicating that the sustained Ca^2+^ mobilizations were dependent upon the presence of nucleotides. Together the data signify that the mobilizations and downstream signals depend upon extracellular ATP (Fig 1C, blue line). Previously, we showed that there was a minor response to EGF and that the EGFR inhibitor, AG1478, suppressed the EGF-induced Ca^2+^ response, but not the ATP-induced response [2]. We also reported that EGFR became phosphorylated on tyrosine residues after injury, and P2Y2 played a role in EGFR cross-activation during cell migration [5, 7]. Therefore, we asked if EGF could induce the sustained Ca^2+^ mobilizations; but did not detect mobilizations above background levels (Fig 1C, yellow line). These findings support the hypothesis that the sustained Ca^2+^ mobilizations are specific to extracellular ATP, indicating that purinoreceptors may play a major role in cell-cell communication.

### Sustained Ca^2+^ mobilizations are mediated through P2X7 and P2Y2 receptors

Through a series of siRNA knockdown and inhibition experiments, we demonstrated that the P2Y2 and P2X7 receptors are major role players in the initial Ca^2+^ response after injury [10-12]. While the cornea expresses a number of P2 purinergic receptors, the latter two receptors have a prominent role in Ca^2+^ mobilization after wounding and cell migration, and their expression changes after injury [7, 10-11, 13, 23]. Given these reports and our observation of sustained extracellular ATP-mediated Ca^2+^ mobilization Fig 1C), we hypothesized that P2X7 and P2Y2 are involved in the sustained Ca^2+^ mobilizations, prompting the development of quantitative methods to examine events of cell communication during the sustained mobilizations. To determine quantitative changes in cell-cell communication, all cells must be inhibited and since siRNA knockdowns were over 60% efficient, we used competitive inhibitors A438079 (for P2X7) and AR-C 118925XX (P2Y2) to achieve a uniform inhibition. The epithelial cells were preincubated with competitive inhibitors to purinergic receptors, wounded, and imaged over time. When cells were incubated with either A438079 (competitive inhibitor to P2X7) (green line) or AR-C 118925XX (competitive inhibitor for P2Y2 receptor) (orange line) and then wounded, the sustained responses were attenuated (Fig 1D).

To analyze the role of the purinoreceptors in cell-cell communication, we examined the sustained Ca^2+^ mobilization patterns in HCLE cells after activation with the agonists, BzATP or UTP, for a minimum of 45 minutes. The concentration of agonist was adapted from receptor kinetics data of the initial wave [21]. We observed that sustained Ca^2+^ mobilizations traveled within groups of three or more cells at any given time, which we defined as a “cluster”, for both P2X7 and P2Y2 receptors (S2 and S3 Movies). The response to UTP was intense and decreased over time, while the response to BzATP had a slower onset and then intensified within clusters of cells (S2 and S3 Movies). We analyzed the Ca^2+^ responses of these clusters with cell-based MATLAB analysis scripts (Fig 2A), which were designed to detect individual cells, and demonstrated that each agonist elicited a unique profile [22]. The analysis revealed that the average percent of active cells and cluster number over time in response to BzATP was less than that detected in response to UTP (Fig 2B). These data indicate that while both agonists generate immediate and sustained Ca^2+^ mobilizations, their output patterns are unique.

**Fig 2.**
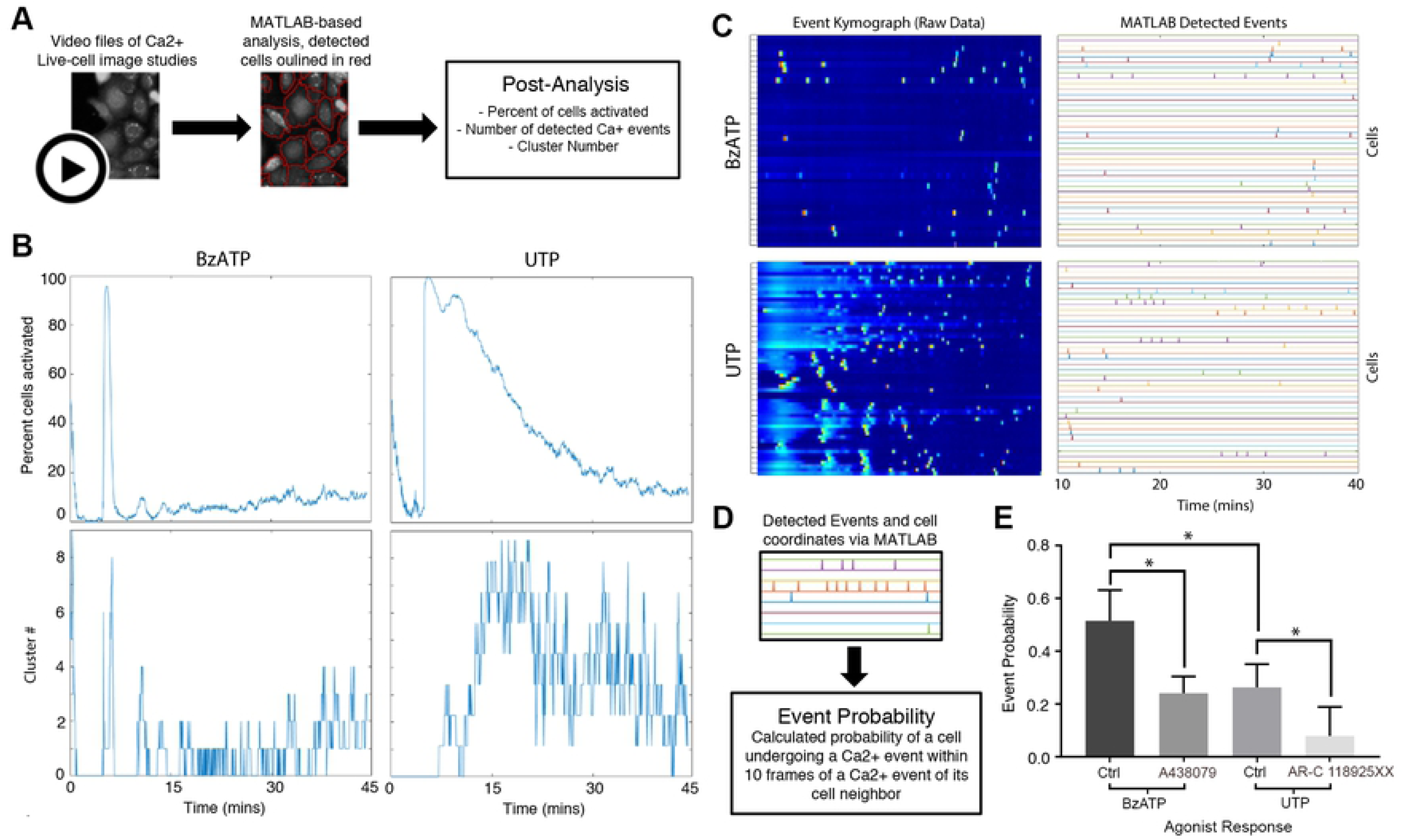
Analysis of UTP and BzATP induced Ca^2+^ mobilizations. (A) Schematic of cell-based approach of the Ca^2+^ analysis. Cells are identified using coordinates from the image study video and event kymographs generated. (B) Representative graphs of percent cells activated over time and cluster number versus time for BzATP and UTP agonist image studies (n=6). (C) Representative kymographs and event charts of UTP and BzATP agonist image studies. To reduce the background noise from the high-intensity initial Ca^2+^ oscillations, mobilizations are analyzed 10 minutes after inducing the Ca^2+^ mobilizations with agonist (n=6). (D) Schematic of event probability based on MATLAB-detected events. (E) Comparison of the average event probability values for each of the agonists and their specific inhibitors, A438079 (P2X7 competitive inhibitor), AR-C 118925XX (P2Y2 competitive inhibitor). Data are means ± SEM and were analyzed with a Tukey’s multiple comparisons test (*p<0.05 for each of the indicated comparisons, n=4).

To quantify the distinct sustained Ca^2+^ mobilization patterns in response to the agonists, kymographs were generated that reflected all of the cells with a known location of each cell. The graph displays activity approximately 10 mins after the immediate Ca^2+^ response (Fig 2C), which allowed for reduction of background noise that occurred due to the high-intensity produced by the immediate Ca^2+^ response. The events that were detected were processed with another MATLAB-based script that calculated “event probability”, which was defined as the probability of one cell displaying a Ca^2+^ event within 10 frames of a detected event of a neighboring cell (Fig 2D). While we demonstrated that BzATP elicited fewer total number of detected Ca^2+^ events compared to UTP (Fig 2C), the average communication event probability for UTP was significantly lower than that for BzATP (*p<0.05) (Fig 2E). These results indicate that the sustained Ca^2+^ mobilization in response to BzATP, while less active overall compared to UTP, exhibits a more coordinated pattern of cell-cell communication. Similar experiments performed with competitive inhibitors to P2X7 or P2Y2 revealed that the Ca^2+^ event probabilities significantly decreased compared to their respective agonist controls (*p<0.05) (Fig 2E). Together these indicate that the receptors most likely to be responsible for cell-cell communication are P2Y2 and P2X7.

While analyzing the event probability for HCLE cells stimulated by an agonist revealed a distinct response, our ultimate goal was to determine the profile of the sustained Ca^2+^ mobilization pattern after injury. Based on our initial observations that the immediate Ca^2+^ response was generated in cells closest to the wound, the cells in wounded culture were categorized into two groups: the first two rows of cells closest to the wound were defined as the leading edge (LE) and the cells in rows further away were defined as back from leading edge (BFLE). The event kymographs and the resulting detected events demonstrated that the LE cells had a larger number of cells exhibiting Ca^2+^ activity compared to BFLE cells (Fig 3A and 3B). When the potential for cell-cell communication was quantified, the average event probability between LE cells was significantly higher (**p<0.01) than that of BFLE cells (Fig 3C). When the LE wounded cell event probability values were compared to the agonist induced events, they were statistically similar to those stimulated with BzATP (Fig 3D). These results imply that the P2X7 receptor may play a role in the healing response of LE cells to coordinate the collective migration process in wound closing. To test the role of P2X7 and P2Y2 in cell-cell communication during wound healing, we calculated the event probability of the LE cells when preincubated with either A438079 or AR-C 118925XX. While the A438079 wounded group had a significantly reduced (***p<0.001) event probability compared to control, the AR-C 118925XX wounded group had no detectable event probability (Fig 3E). While the wound response after pretreatment with AR-C 118925XX did have visible Ca^2+^ events, they were not between neighboring cells, which is required to calculate the event probability values. Given that both receptors are activated by the ATP released from wounded epithelial cells, these differing results imply that Ca^2+^ signaling is orchestrated via cooperation between P2X7 and P2Y2 receptors.

**Fig 3.**
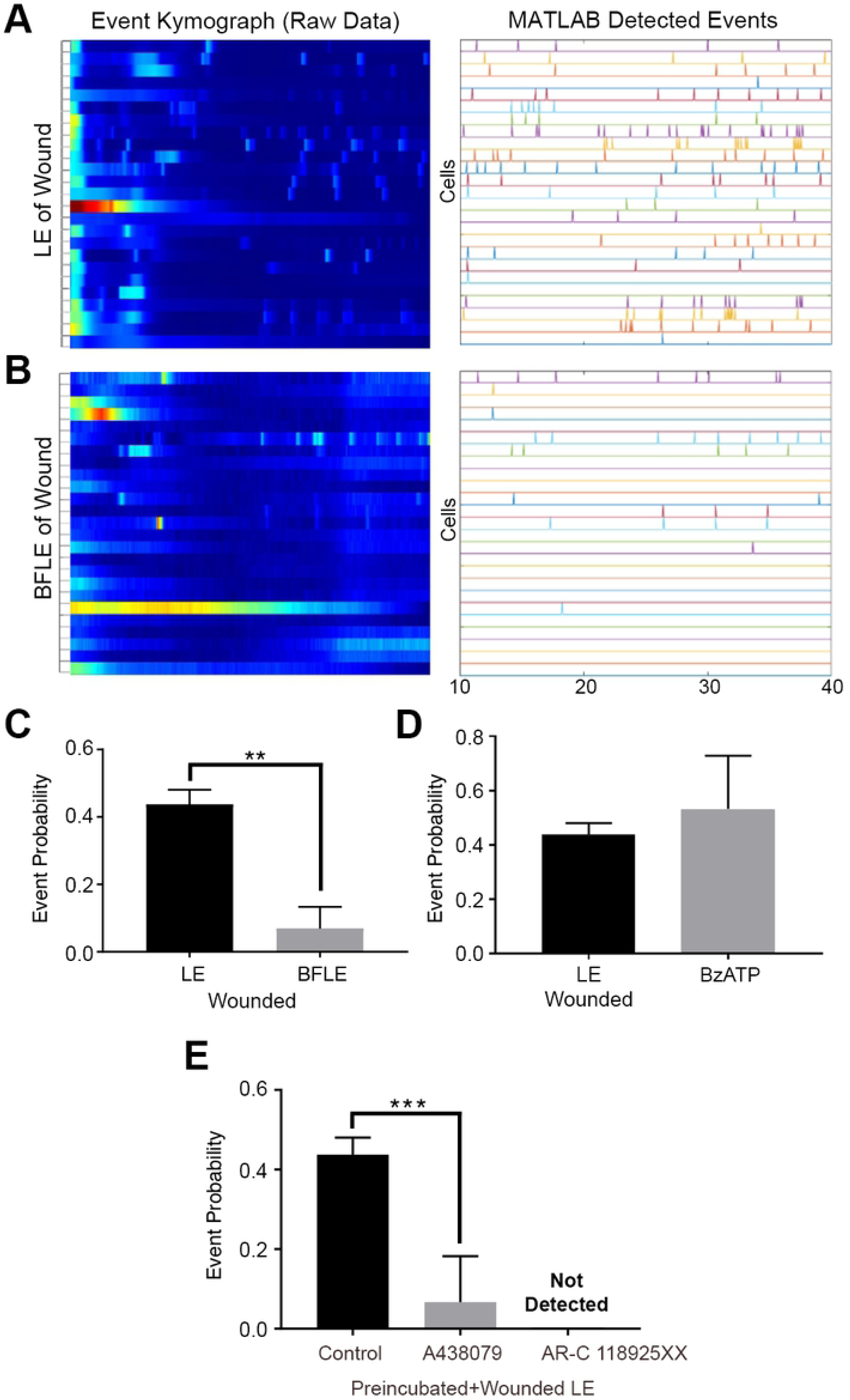
Communication events between cells depend on distance from wound. (A and B) Representative kymographs and detected event charts of the leading edge (LE) (A) and back from the leading edge (BFLE) (B). Analysis was performed 10 minutes after wounding. (n=3). (C) Event probability values for LE and BFLE cells after wounding. Data are mean ± SEM and were analyzed with a two-tailed unpaired t-test (**p<0.002, n=3). (D) Event probability values for the LE wound and BzATP agonist response. Data are mean ± SEM and were analyzed with a two-tailed unpaired t-test (ns, n=3). (E) Event probability values for the LE when cells were preincubated in the presence or absence of A438079 or AR-C 118925XX before scratch-wounding. Data are mean ± SEM and were analyzed with a one-way ANOVA with the Tukey’s multiple comparisons test (***p<0.001, n=4).

### Activation of purinoreceptors promote cell migration after injury

To examine the role of the purinoreceptors on cell motility, cells were loaded with Fluo-3AM (cyan) and CellMask™ (Fire LUT), injured, and monitored over several hours (Fig 4A, S4 Movie). Cells either displaying sustained Ca^2+^ mobilizations or lack thereof were classified as “active” and “inactive” cells respectively. The two groups of cells were tracked with CellMask™ membrane dye, and the cell membrane traces were used to record motility and change in cell shape (Fig 4B). Active cells exhibited a change in cell shape over time and demonstrated cell motility (Fig 4A). Based on these observations, we hypothesized that sustained Ca^2+^ mobilization patterns, altered cellular morphology and motility are necessary for proper wound healing, and these events play a role in orchestrating collective epithelial cell migration during wound repair.

**Fig 4.**
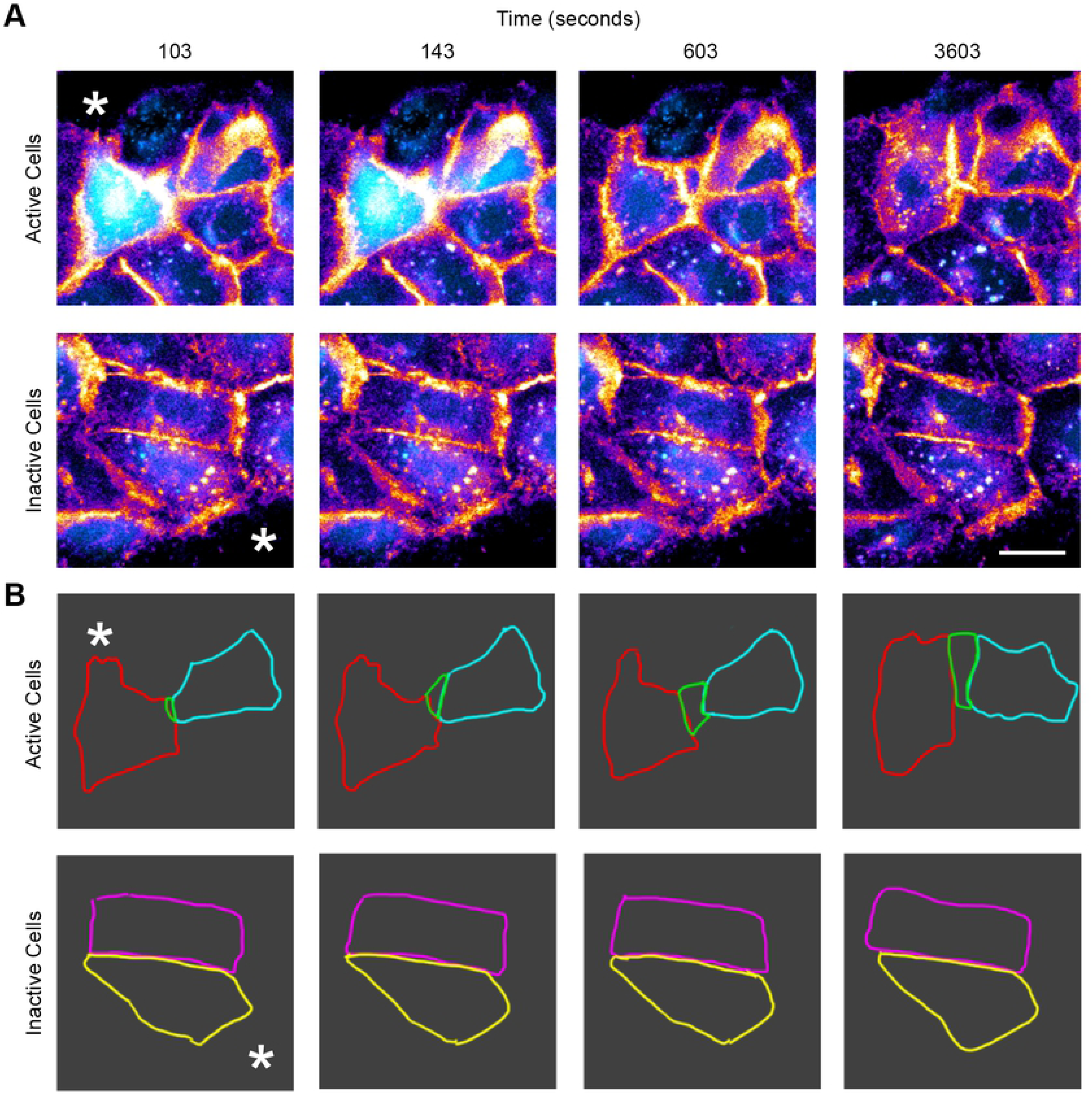
Ca^2+^ mobilizations between cells correlate with changes in cell shape. (A) Live-cell imaging of the wound edge. Cells were incubated with CellMask™ membrane dye (Fire LUT) and Fluo-3AM (cyan) 10 minutes after injury to examine cell shape changes and Ca^2+^ mobilizations. The edge of the wound is marked with an asterisk. (B) Cell traces were made for cells that exhibited active and inactive Ca^2+^ mobilization. When Ca^2+^ mobilizations are present between cells (Active), changes in cell shape are detected. When Ca^2+^ mobilizations are absent (Inactive), there is no detectable change in cell morphology. Scale bar = 34 μm. (n=3 for both A and B).

### Ca^2+^ mobilizations between cells occur through pannexin channels but not connexin gap junctions

In order to determine how sustained Ca^2+^ mobilizations transmitted from cell-cell, we examined the role of connexin gap junctions, specifically connexin 43 (Cx43), and pannexin channels in cell-cell communication. Immunohistochemistry studies demonstrated that Cx43 was present as punctate staining (Fig 5A, yellow) along the cellular membrane in HCLE cells seeded at a high density, but not at a low density (Fig 5A). To test whether gap junctions were responsible for the transmission of sustained Ca^2+^ mobilizations, we preincubated the cells with alpha-glycyrrhetinic acid (α-GA), a connexin-specific inhibitor that disassembles junctions [2]. Utilizing the cell-based MATLAB analysis scripts, we demonstrated that while α-GA dampens the % of activated cells, it does not alter the average cluster number (Fig 5B). Furthermore, there was no significant difference in the mean event probability values between the two groups (Fig 5C). These results indicate that propagation of events between cells does not occur via gap junctions but instead through some other means.

**Fig 5.**
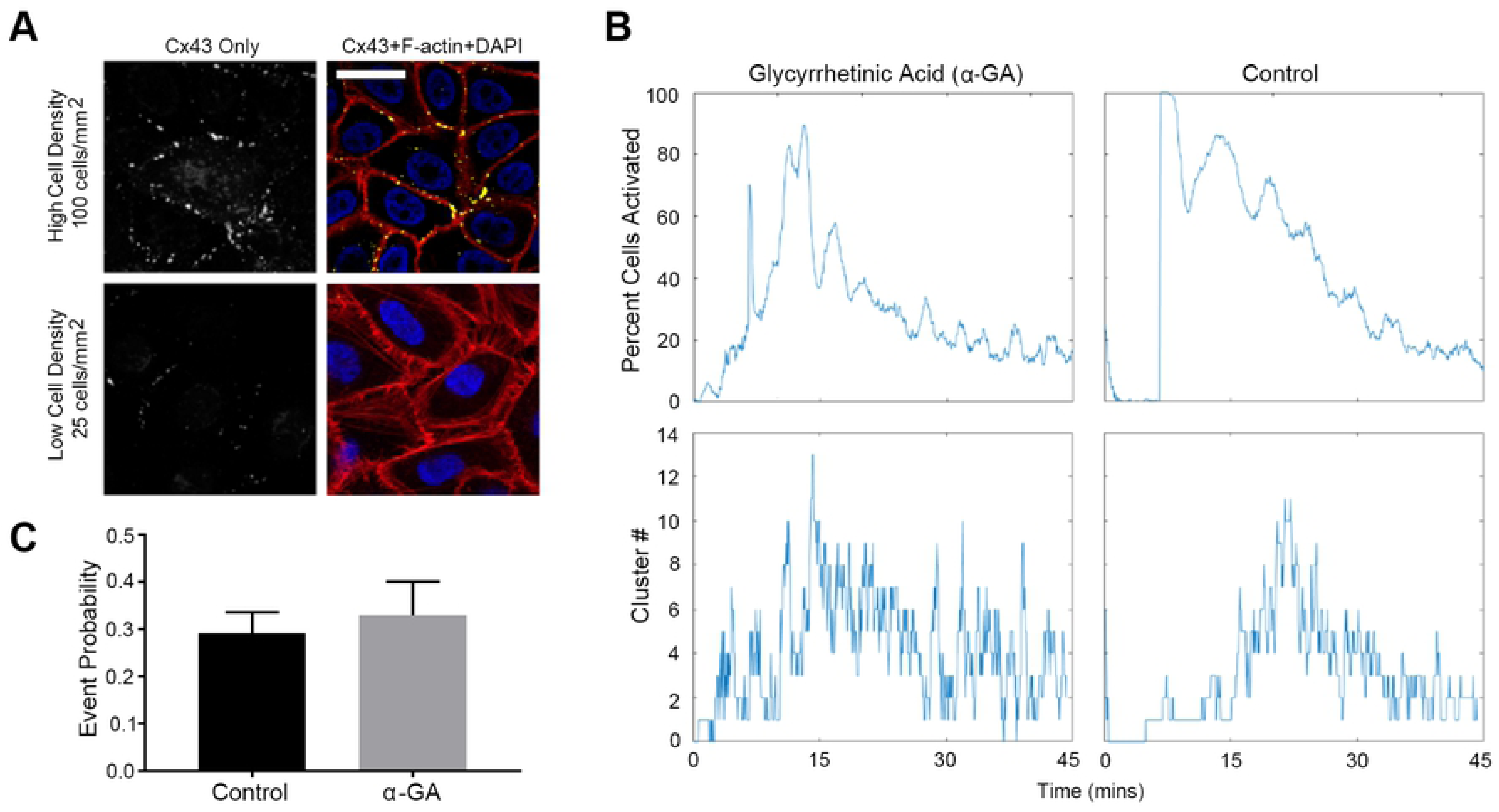
Connexin-43 (Cx43) channels do not mediate the mean event probability. (A) Localization of Cx43 (white in the Cx43 only image, yellow in the composite image). Cells were counter-stained with rhodamine phalloidin (red) and DAPI (blue). Higher cell density and confluence correlated with extend of localization of connexin along the cell membrane. Scale bar = 42 μm. (n=4). (B) Representative graphs of percent cells activated over time and cluster number versus time for Ca^2+^ mobilization determined from videos of cells preincubated with 120 μM α-GA and control. (n=4). (C) Event probability values for the control and α-GA treated group. Data are mean ± SEM and were analyzed with a two-tailed unpaired t-test (ns, n=3).

A second candidate communication pathway is pannexin, specifically pannexin 1. Our previous observation that apyrase quenched the Ca^2+^ response led us to hypothesize that pannexin1’s localized ATP release was responsible for the propagation of the sustained Ca^2+^ mobilizations. To test whether inhibiting pannexin would affect Ca^2+^ mobilizations, we used 10Panx, a pannexin-specific inhibitor, and the scramble Panx peptide control (Ctrl) [24-25]. When cells were preincubated with 10Panx and stimulated with BzATP, there was a significant decrease in the percent of activated cells and cluster number over time in the 10Panx group compared to Ctrl (Fig 6A). We also demonstrated that P2X7 interacted with pannexin1 in epithelial cells using in situ crosslinking studies (S2 Fig). Stimulation of cells using the agonists, BzATP and UTP, allowed us to demonstrate that inhibition of pannexin channels abrogated the sustained Ca^2+^ mobilizations stimulated with BzATP. Furthermore, inhibition with 10Panx, resulted in an event probability that was significantly decreased (**p<0.009) (Fig 6B); whereas stimulation of cells previously inhibited with 10Panx stimulation did not significantly reduce cell-cell communication (Fig 6B). Together these data indicate the participation of pannexin channels in Ca^2+^ mobilization.

**Fig 6.**
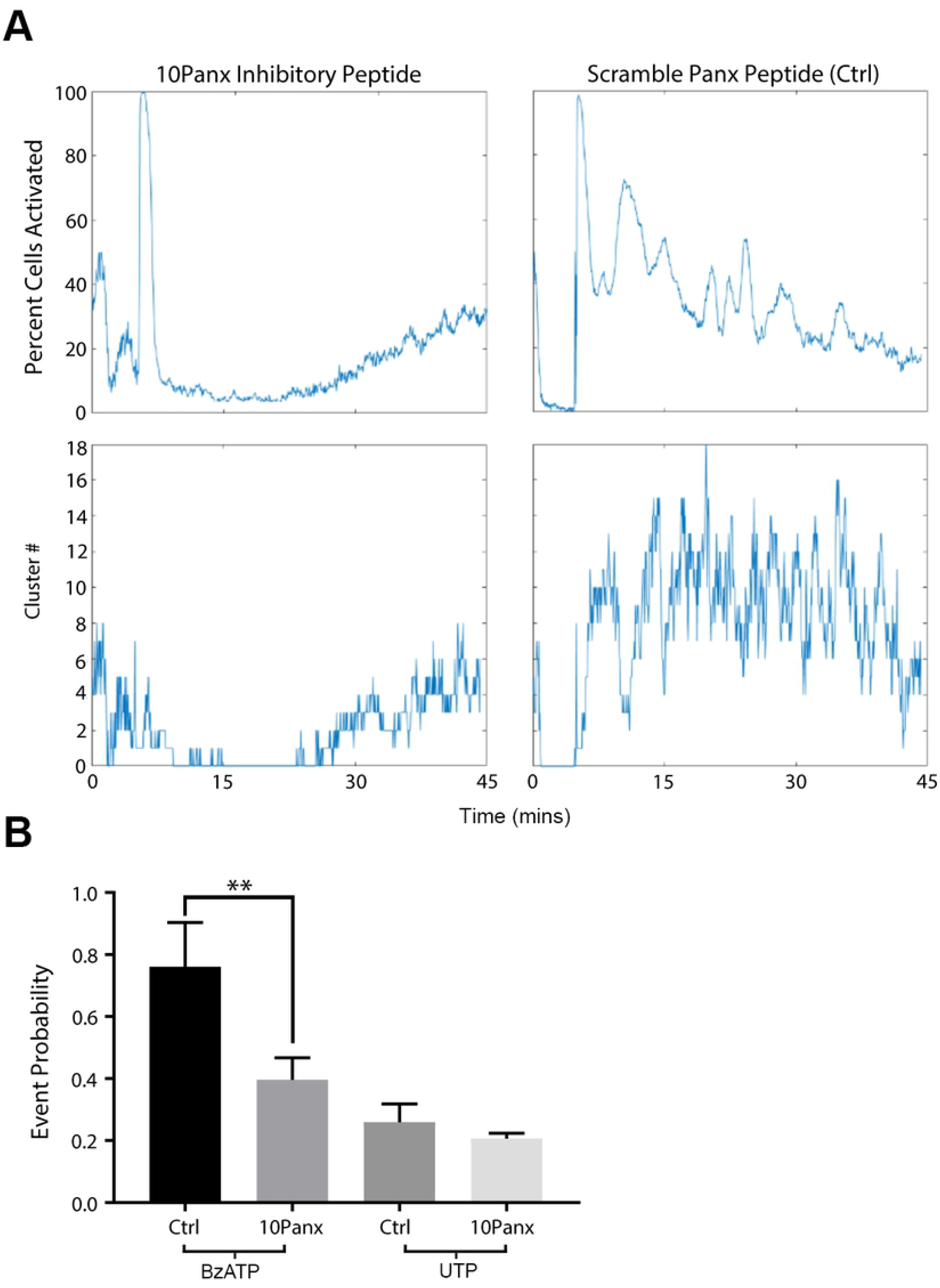
Pannexin1 facilitates the propagation of Ca^2+^ mobilizations when purinergic receptors are activated. (A) Representative graphs of percent cells activated over time and cluster number when cells were preincubated with 100 μM 10Panx inhibitory peptide or scrambled peptide control prior to stimulation. (n=4). (B) Event probability values of cells preincubated with either 10Panx or Scrambled Panx control group and activated with either BzATP or UTP. Data are mean ± SEM and were analyzed with a two-tailed unpaired t-test. 10Panx significantly lowered cell-cell communication if cells were stimulated with P2X7 (**p<0.009), while communication was unaffected when stimulated with UTP (ns, n=4).

To understand the role of ion channels in wound healing, the localization of pannexin1 (Fig 7A, B; yellow) was examined in control and wounded conditions in vitro and in tissue. Pannexin1 was localized at the intercellular space of confluent unwounded epithelial cells in culture and 30 minutes after wounding (Fig 7A; * indicates wound), it was detected also at the leading edge of the wound. However, by two hours it was prominent along the wound (arrows). In corresponding unwounded mouse corneal tissue (Fig 7B; arrowheads), pannexin1 localization was similar to the confluent cells (Fig 7A. Within two hours after wounding, pannexin1 was punctate and present for several cells back from the leading edge of the wound, and by four hours the localization was prominent (Fig 7B, arrows). This change in pannexin1 localization may explain why the sustained Ca^2+^ mobilizations after injury were present predominantly in cells closest to the wound edge (Figs. 1,3). Therefore, cells were incubated in the presence or absence of 10Panx prior to a scratch wound and the event probability analyses were applied to the videos and cells at the LE were analyzed (Fig 7C, D). Cells treated with 10Panx had fewer detected Ca^2+^ events compared to Ctrl (Fig 7C), resulting in significantly lower average event probability values in the 10Panx-treated group (Fig 7D; **p<0.01). These results led us to hypothesize that pannexin inhibition would also affect cell migration and wound closure. To study this, we used long-term live cell imaging of cells preincubated with SiR actin Spirochrome to examine cell migration and wound closure in the presence of 10Panx and scrambled control peptide (Fig 8). The cell traces of the epithelial cells obtained from the migration videos demonstrated that 10Panx inhibited wound closure rate and altered cell migration (Fig 8A, B, C; S5 and S6 Movies). The individual cells revealed different trajectories at the LE compared to those BFLE (S5 and S6 Movies). As shown in Figure 8B, the rate of closure was initially faster in the 10Panx group (red) but over time, the control group’s wound closure rate increased while the inhibitor group stagnated, resulting in delayed wound closure for the 10Panx group (Fig 8B). The individual trajectories were analyzed and the data was organized and presented as two cell groups, LE and BFLE cells (Fig 8C). The LE cells in both groups generally moved in the direction of the wound (Fig 8C), while the BFLE cells in the control wounds moved in the direction of the wound, the majority of the cells pretreated with 10Panx did not move in the forward direction and instead moved laterally. These findings support our hypothesis that pannexin channels are crucial players in sustained Ca^2+^ mobilization and cell migration in corneal epithelial cells.

**Fig 7.**
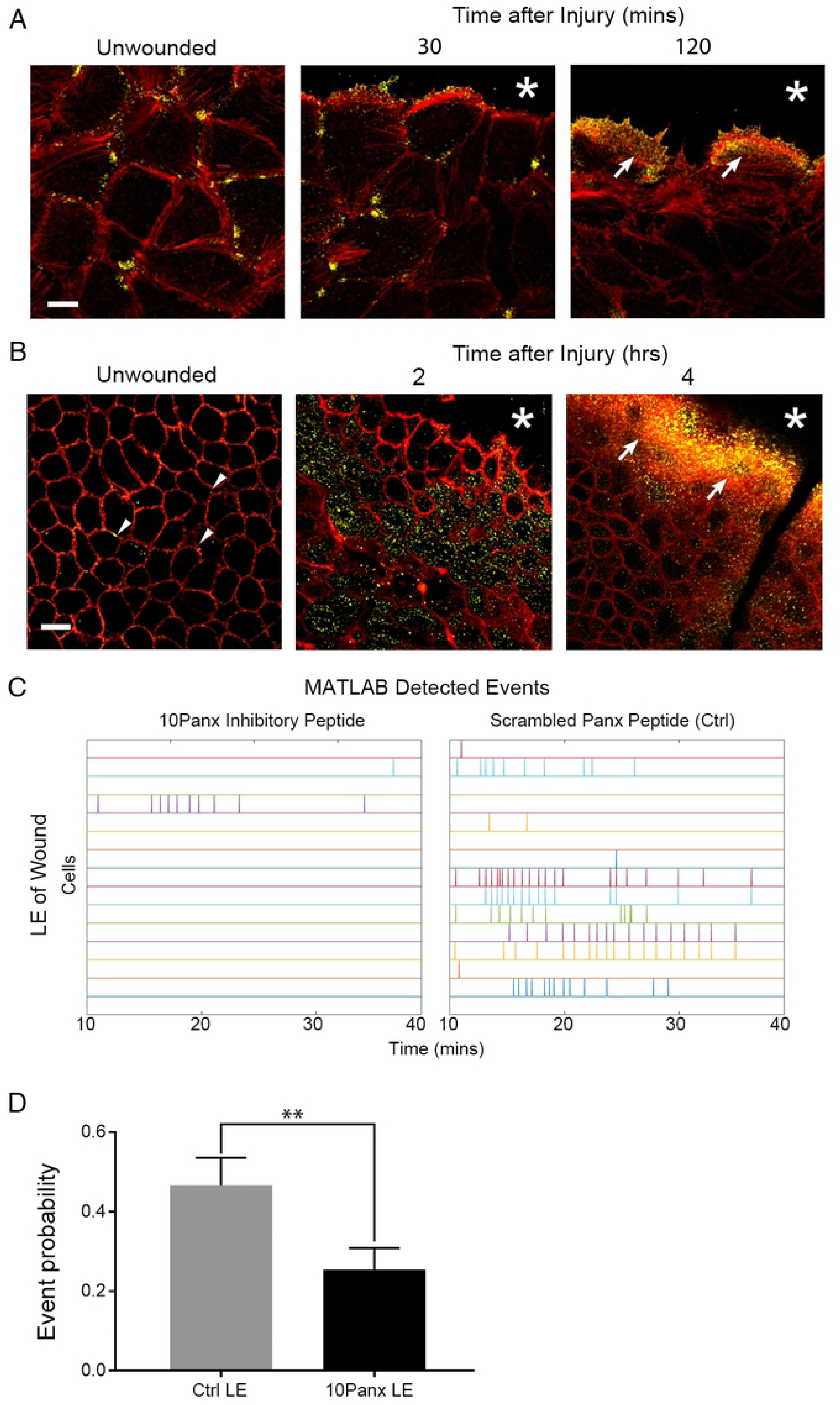
Pannexin1 localization is detected at wound edge during healing. (A) Representative confocal immunofluorescence images of cultured cells stained for pannexin1 localization (yellow, including arrowheads) and counterstained with rhodamine phalloidin (red). Pannexin1 is concentrated adjacent to the leading edge of the wound. Scale bar = 23 μm. (n=3). (B) Representative confocal immunofluorescence images of basal corneal epithelium stained for pannexin1 (yellow, including arrows) and counterstained for rhodamine phalloidin (red). After wounding (2 and 4 hours) the pannexin1 concentrated towards the leading edge of the wound. An asterisk indicates the leading edge of migrating epithelium. Scale bar = 18.5 μm. (n=3). (C) Ca^2+^ mobilizations are represented over time in event charts of LE and BFLE cells 10 minutes after wounding, with cells preincubated with inhibitory peptide 10Panx or scrambled peptide control (Ctrl) (n=3). (D) Event probability values for LE cells after wounding for both treated and Ctrl groups. Data are mean ± SEM and were analyzed with a one-way ANOVA with the Tukey’s multiple comparisons test (**p<0.01, n=3).

**Fig 8.**
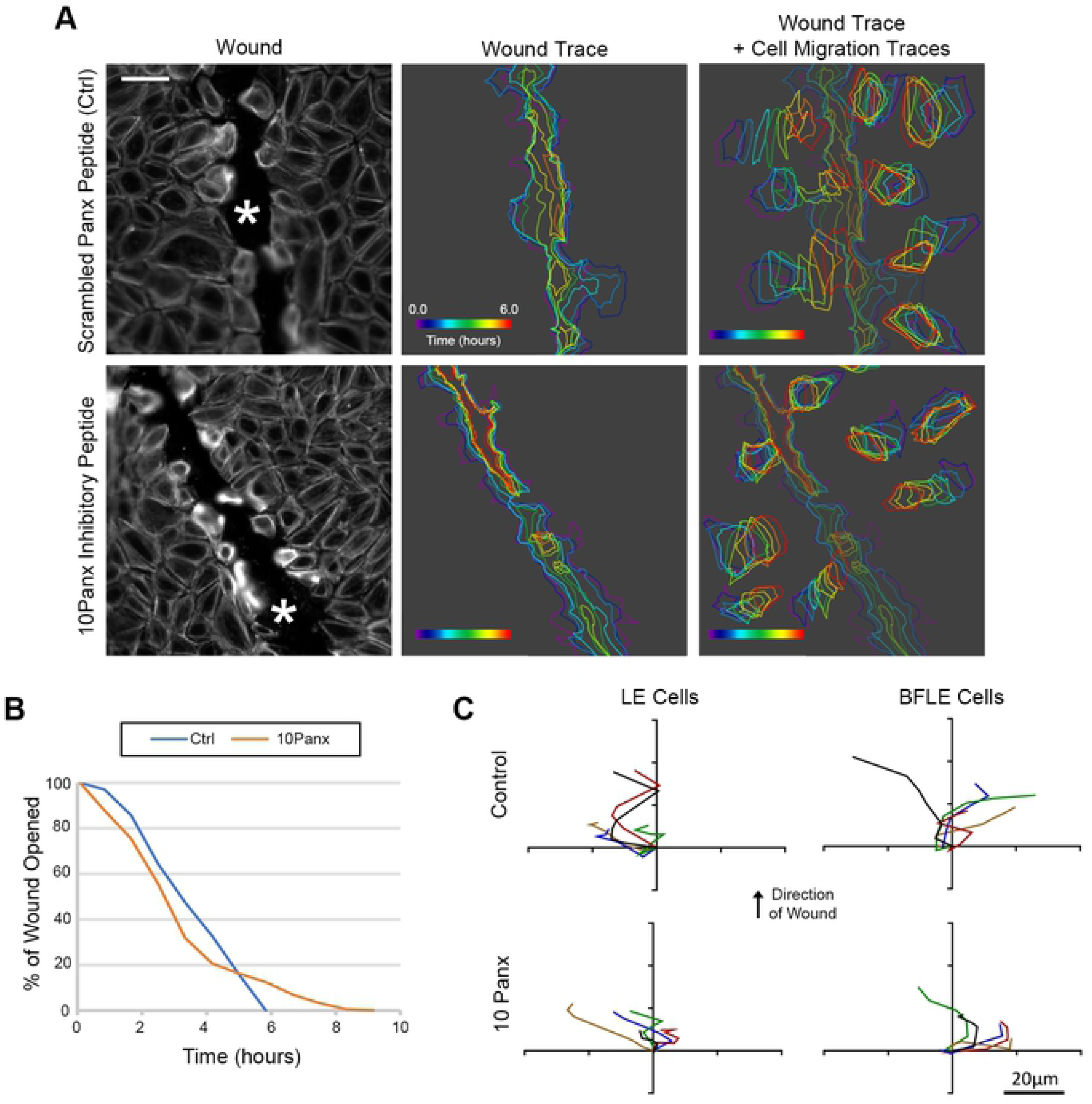
Inhibition of pannexin channels affect cell migration and wound healing. (A) Live-cell imaging of the wound edge of cells preincubated with SiR-Actin Spirochrome (in grayscale) to determine cell migration over a 16 hour period. Scale bar = 66 μm. Traces of the wound area and 8 random cells in the field were drawn over time to observe the rate of cell migration and wound closure for the experimental and control conditions. Colors reflect time and are indicated by time wedge (n=3). (B) Representative percent wound closure graph of cells preincubated with 10Panx or Panx scrambled peptide control over time (n=3). (C) Representative cell migration trajectory diagrams of of LE and BFLE cells preincubated in either 10Panx or Panx scrambled peptide control. Each line represents the migration path of a single cell plotted from a common origin. Scale bar = 20 μm.

### Sustained Ca^2+^ mobilizations are detected in ex vivo models of the cornea

Previously, work on Ca^2+^ mobilizations has been performed primarily on in vitro corneal models [2, 10, 21]. The next logical step is to confirm the presence of the sustained mobilizations in animal models. Therefore, we examined if treatment with an agonist would induce a sustained Ca^2+^ response in the mouse cornea. Live-imaging performed after the eyes were preincubated in Fluo-3AM and CellMask™ (See Methods). Use of the CellMask™ (red) allowed for imaging of specific layers of cells. Images are not displayed for the first 10 minutes after the stimulation because of the noise generated by the initial transient wave as described previously(Fig 1). When corneal epithelial cells were stimulated with ATP, the basal cells exhibited sustained Ca^2+^ mobilizations (Fig 9A, S7 Movie) and these events were examined using MATLAB analyses to generate the event kymograph and detected events (Fig 9B). The results demonstrate that it is feasible in the future to apply our image analyses on corneas with different pathologies and conditions.

**Fig 9.**
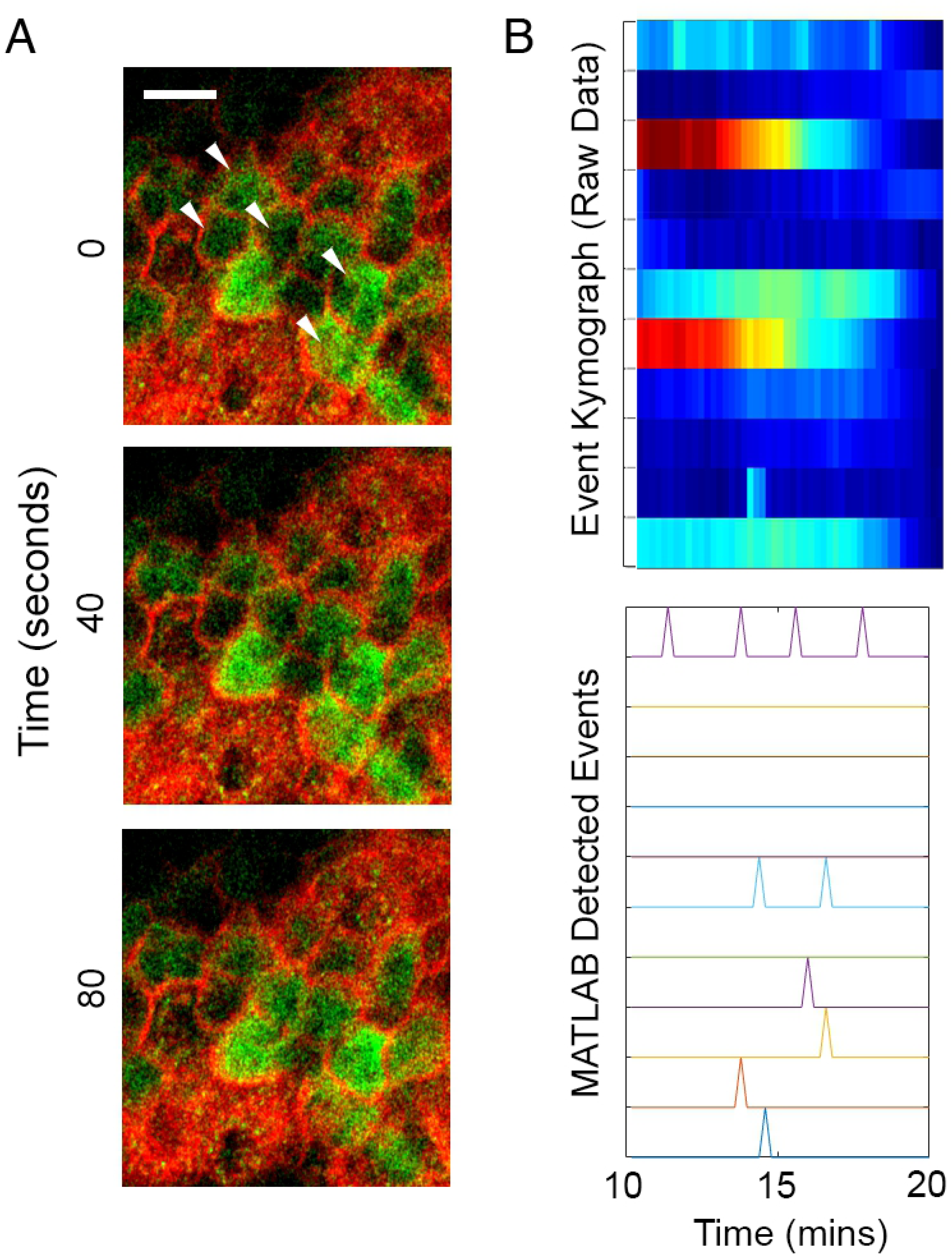
Ca^2+^ mobilizations are present in ex vivo organ cultures after stimulation with ATP. (A) Representative images of Ca^2+^ mobilizations in mouse corneal epithelium (white arrowheads) Similar to in vitro experiments, images were not recorded until 10 minutes after addition of agonist. Scale bar = 15 μm. (n=2). (B) Representative kymograph of Ca^2+^ mobilizations and graph representing detected events after stimulation.

## Discussion

While the importance of epithelial sheet migration is well recognized, there is a lack of understanding in the role of communication that occurs between cells after injury [26]. One way of examining how epithelial cells move is to evaluate the dynamic communication that occurs after injury and understand how it is coordinated and this can be examined by evaluating the role of Ca^2+^ signaling in orchestrating cell migration and wound repair.

This study examined the role of the sustained Ca^2+^ mobilizations that were generated either after an epithelial injury or treatment with an agonist, and that lasted for a period of several hours. Previously we showed that an initial transient response occurred with injury and proposed that it mediated downstream signaling events by altering phosphorylation of focal adhesion and adaptor proteins in a phosphoproteomic study [2, 7]. Furthermore, knocking down certain purinergic receptors confirmed that this family of receptors did mediate the response [7]. In our current experiments we demonstrated that neighboring cells display synchronous mobilizations at the wound edge, which decreased over time and distance from the wound edge. Sustained Ca^2+^ mobilizations are not limited to injury and have been reported in a number of developmental systems. These events or mobilizations were hypothesized to guide cell migration in zebrafish and modulate changes in IP3- mediated Ca^2+^ release from an oscillatory to a tonic mode [27-28]. In addition, they were detected during status epilepticus where Ca^2+^ waves continue for extended time periods [29]. Furthermore, still other investigators proposed that short flickers of Ca^2+^ may mediate the directionality of cell migration [30].

To examine the response, we used several approaches to analyze Ca^2+^ mobilization and cell communication. Experiments where apyrase, an ectonucleotidase, was added prior to the sustained Ca^2+^ mobilizations demonstrated that extracellular ATP was required. Since this Ca^2+^ response required extracellular ATP, we examined the potential role of purinoreceptors in the sustained response as corneal epithelial cells express P2X7 and P2Y2 and both have been shown to play a role in cell migration [7, 10]. The sustained injury-induced response was abrogated in the presence of competitive inhibitors to these receptors. While it is possible that the EGF receptor may mediate these features of cell-communication, it in of itself had a negligible effect. The differences in communication, frequency and intensity are similar to events found in development [31]. These similarities are not unexpected as wound repair or directed migration after an injury may have a number of stimuli that are similar to developing systems.

To quantify the response, we developed image processing techniques to monitor the cells and examine their interactions through percent of active cells, cluster number, and probability event values. These tools were developed to analyze the response to agonist stimulation and then applied to the wound response. For example a qualitative assessment of the response to the agonist UTP indicated that the majority of the cells appeared to be in an “on or off state”, while in response to BzATP there were regions of high activity and regions of low activity. A quantitative analysis verified that the Ca^2+^ response to BzATP elicited a lower percent of active cells and cluster number compared to UTP. However, the sustained Ca^2+^ mobilization in response to BzATP, while less active overall compared to UTP, exhibited a more coordinated pattern of cell-cell communication as demonstrated by higher event probabilities. Additional experiments performed with competitive inhibitors of the P2X7 or P2Y2 receptors revealed that the Ca^2+^ event probabilities decreased compared to their respective agonist controls. These indicated that the receptor most likely responsible for cell-cell communication was the P2X7 receptor.

We provide evidence that the sustained Ca^2+^ mobilizations along the wound edge between cells are critical for the onset of cell motility. Mobilizations between cells near the wound edge were correlated with a change in cell shape when cells were co-stained with CellMask™ and Fluo-3AM and these were supported by analyses revealing significant differences in event probabilities between the LE and BFLE groups, with LE cells having higher probability values than the BFLE cells. Interestingly, the event probability values of LE cells were similar to those when cells were stimulated with BzATP, suggesting that the P2X7 receptor may be involved in the wound healing response induced by the LE cells. This response could explain how injured cells coordinate themselves for collective migration to close the wound and is supported by studies demonstrating the transient localization of P2X7 at the wound edge [10].

The coordinated activity in the Ca^2+^ response to BzATP may be explained in part by the fact that the P2X7 receptor is a channel that allows for ATP transport in and out of cells, resulting in a positive feedback by allowing the cells at the leading edge to function as mechanically coupled yet electrochemically isolated units [32]. Preliminary experiments revealed that thapsigargin, an IP3 mediated inhibitor, diminished ruffling at the edge and these were associated with a change in Ca^2+^ mobilization [20]. Evidence from other cell systems suggests the presence of a feed-forward system where ATP could move through pannexin channels and activate P2X7 receptors [14]. This suggests that there is a continuous release of ATP along the wound margin developing a chemotactic gradient for the migrating cells that is associated with the sustained Ca^2+^ mobilizations. Previously investigators have demonstrated that the ATP released by neutrophils acts as a chemoattractant [15-17].

Although the activity of the sustained Ca^2+^ mobilizations is cell density dependent, the probability that cell-cell communication propagated through gap junctions was not reduced with alpha-glycyrrhetinic acid, a specific inhibitor of gap junctions, that disrupts the junctions. Another candidate channel protein, pannexin, may be the more likely candidate [3, 24]. Its role is demonstrated in dendritic cells pannexin1 and P2X7 where both proteins play a role in cell migration during injury [18]. Using a specific pannexin channel inhibitor, we demonstrated that cell migration rate, cell behavior during migration and Ca^2+^ mobilization were altered when pannexin1 was inhibited. Studies where communication or event probability was assessed after cells were incubated in the presence or absence of 10Panx and then stimulated with UTP or BzATP revealed that the probability of communication was impeded significantly when cells were activated with BzATP. Our current proposed model for Ca^2+^ mobilization propagation is localized release of ATP through pannexin channels activating purinergic receptors in neighboring epithelial cells. Specifically in our epithelial cells, ATP remained 6-to 7-fold higher after injury compared to the near constant basal levels of unwounded control cells. These indicate that there may be an overall greater release than degradation of ATP as it may be secreted constantly by migrating cells. These concur with the observation of cells at the leading edge where mobilization of Ca^2+^ was associated with rapid changes in cell morphology and migration.

Study of the communication between cells provides insight into the mechanisms of wound repair in control and diseased conditions. The epithelial injury model and the quantitative processing provides a valuable system to investigate how cells communicate in response to specific receptors. This model can be used to identify therapeutic targets and test strategies in the cornea and in other tissues to modulate the collective cell migration in treating and preventing disease progression.

## Acknowledgements

We thank Drs. Brigitte Ritter, Dr. Matthew Nugent, and Gregory Teicher for critical and challenging discussions. We thank Ms Audrey Hutcheon for editorial comments.

## Supporting information

**S1 Fig Representative kymograph of cells at least 2 cell rows away from the wound edge.** Compared to the kymographs made from cells at the wound edge (LE), the Ca^2+^ response showed less intensity. Brackets on the left and each horizontal line represent activity of a single cell (n=7).

**S2 Fig Association of P2X7 and pannexin1 protein in epithelial cells**. HCLE cells were cultured until confluent, and cross-linking was performed with formaldehyde in situ, as previously described [12]. Each crosslinked experimental sample (labeled “CL”) and its corresponding control were heated at two different temperature settings: 65°C (to maintain crosslinks) and 95°C (to disrupt crosslinks). Both CL lanes displayed the crosslinked P2X7+ pannexin1 protein product, with the CL (95°C) lane verifying the composition crosslinked protein product. (n=3).

**S1 Movie. Sustained Ca**^**2+**^ **oscillations detected after scratch-wounding.** Confluent cells were preincubated with 5 μM of Fluo3-AM for 30 minutes. Cells were scratch-wounded and imaged for 2 hours in an environmental chamber mounted on a Zeiss 880 confocal microscope (10x). Images were taken every 3 seconds, with the movie at 25 fps. Scale bar = 60m.

**S2 Movie. Sustained Ca**^**2+**^ **oscillations induced by UTP.** Confluent HCLE cells were preincubated with 5μM of Fluo3-AM for 30 minutes. Cells were stimulated with 25 μM UTP and imaged for 45 minutes in an environmental chamber mounted on a Zeiss 880 confocal microscope (20x). Images were taken every 3 seconds, with the movie at 25 fps. Scale Bar = 50 μm.

**S3 Movie. Sustained Ca**^**2+**^ **oscillations induced by BzATP stimulation.** Confluent HCLE cells were preincubated with 5 μM of Fluo3-AM for 30 minutes. Cells were stimulated with 25 μM BzATP and imaged for 45 minutes in an environmental chamber mounted on a Zeiss 880 confocal microscope (20x). Images were taken every 3 seconds, with the movie at 25 fps. Scale Bar = 50 μm.

**S4 Movie. Ca**^**2+**^ **mobilizations and cell shape.** Confluent HCLE cells were preincubated with 5 μM Fluo3-AM for 30 minutes and CellMask™ Deep Red Plasma membrane stain at recommended concentration for 5 minutes. Cells were scratch-wounded and imaged for 45 minutes in an environmental chamber mounted on a Zeiss 880 confocal microscope (40x oil). Images were taken every 5 seconds, with the movie at 25 fps. Scale Bar = 34 μm.

**S5 Movie. 10Panx significantly attenuates wound closure rate.** Confluent HCLE cells were treated with 100 μM 10Panx inhibitory peptide for an hour before being preincubated with 5 μM Fluo3-AM for 30 minutes. Cells were scratch-wounded and imaged for 16 hours in an environmental chamber mounted on a Zeiss 880 confocal microscope (20x). Images were taken every 5 minutes, with the movie at 50 fps. Scale Bar = 66 μm.

**S6 Movie. Pannexin scrambled peptide does not inhibit rate of wound closure.** Confluent cells were treated with 100 μM Scrambled Panx control peptide for an hour before being preincubated with 5 μM Fluo3-AM for 30 minutes. Cells were scratch-wounded and imaged for 16 hours in an environmental chamber mounted on a Zeiss 880 confocal microscope (20x). Images were taken every 5 minutes, with the movie at 50 fps. Scale Bar = 66 μm.

**S7 Movie. Ca**^**2+**^ **mobilizations in organ culture.** Mouse corneas were preincubated with 15 μM Fluo3-AM for 30 minutes and CellMask™ Deep Red Plasma membrane stain at recommended concentration for 5 minutes. Cells were scratch-wounded and imaged for at least 15 mins in an environmental chamber mounted on a Zeiss 880 confocal microscope with AIRYSCAN Fast Module (20x). Images were taken every 10 seconds, with the movie at 25fps. Scale Bar = 16.5 μm.

